# Characterisation of *Medicago truncatula* CLE34 and CLE35 in nodulation control

**DOI:** 10.1101/2020.07.31.231605

**Authors:** Celine Mens, April H. Hastwell, Huanan Su, Peter M. Gresshoff, Ulrike Mathesius, Brett J. Ferguson

## Abstract

Legume plants form a symbiosis with N_2_-fixing soil rhizobia, resulting in new root organs called nodules that enable N_2_-fixation. Nodulation is a costly process that is tightly regulated by the host through Autoregulation of Nodulation (AON) and nitrate-dependent regulation of nodulation. Both pathways require legume-specific CLAVATA/ESR-related (CLE) peptides. Nitrogen-induced nodulation-suppressing CLE peptides have not previously been characterised in *Medicago truncatula*, with only rhizobia-induced MtCLE12 and MtCLE13 identified. Here, we report on novel peptides MtCLE34 and MtCLE35 in nodulation control pathways. The nodulation-suppressing CLE peptides of five legume species were classified into three clades based on sequence homology and phylogeny. This approached identified MtCLE34 and MtCLE35 and four new CLE peptide orthologues of *Pisum sativum*. Whereas MtCLE12 and MtCLE13 are induced by rhizobia, MtCLE34 and MtCLE35 respond to both rhizobia and nitrate. MtCLE34 was identified as a pseudogene lacking a functional CLE-domain. Overexpression of *MtCLE12, MtCLE13* and *MtCLE35* inhibits nodulation. Together, our findings indicate that MtCLE12 and MtCLE13 have a distinct role in AON, while MtCLE35 regulates nodule numbers in a rhizobia- and nitrate-dependent manner. MtCLE34 likely had a similar role to MtCLE35 but its function was lost due to a nonsense mutation resulting in the loss of the mature peptide.

## Introduction

Nitrogen availability is essential for plant growth, but is highly variable in the rhizosphere with up to 1,000-fold differences in nitrate concentration (μM-mM) reported within a four-meter diameter (Miller *et al*., 2007). Plants encountering nitrogen deficiency require an increase in root plasticity to maintain an optimal nitrogen balance (*i*.*e*., via lateral root growth, increased uptake or soil microbial interactions). Legume plants have established a symbiotic relationship with rhizobia bacteria that fix atmospheric nitrogen (N_2_) for plant-use in return for photosynthetic carbohydrates. This requires the formation of specialised root nodules that house the bacteria. Biological nitrogen fixation and nodule organogenesis are both resource intensive processes and thus the host plant has evolved regulatory mechanisms to control nodule numbers, such as the Autoregulation of Nodulation (AON) pathway (Caetano-Anollés & Gresshoff, 1991a, 1991b; Delves *et al*., 1986; Ferguson *et al*., 2019).

AON is a systemic feedback mechanism requiring extensive signalling between the roots and shoot Moreover, when ample nitrogen is present in the soil, nodulation is negatively controlled via nitrate-dependent regulation of nodulation which has many signals in common with AON. In fact, the first AON mutants were identified as nitrate-tolerant supernodulators (*nts*) (Carroll *et al*., 1985). Both processes initially require upregulation of CLAVATA3/EMBRYO SURROUNDING REGION-RELATED (CLE) peptides that are perceived by a leucine-rich repeat receptor-like kinase (LRR-RLK) called MtSUNN/GmNARK/LjHAR1/PsSYM29/PvNARK (Krusell *et al*., 2002; Nishimura *et al*., 2002; Searle *et al*., 2003; Schnabel *et al*., 2005; Ferguson *et al*., 2014). The shoot-derived signal of AON has been identified as miR2111, which targets the Kelch-repeat factor, TOO MUCH LOVE (TML), for degradation to inhibit the development of new nodules (Takahara *et al*., 2013; Tsikou *et al*., 2018; Gautrat *et al*., 2020). Recent findings using *Lotus japonicus* suggest TML might not be required for nitrate-dependent inhibition (Nishida *et al*., 2020).

CLE peptides are hormone-like regulators of cell division and differentiation acting in local and systemic cell-to-cell communication. Mature CLE peptides are 12 to 13 amino acids in length and are post-transcriptionally processed from the C-terminal region of their respective pre-propeptides (Hastwell *et al*., 2015b). They require further modification through triarabinosylation of a central hydroxylated proline residue by members of the hydroxyproline *O*-arabinosyltransferase family (Okamoto *et al*., 2013; Corcilius *et al*., 2017; Kassaw *et al*., 2017; Hastwell *et al*., 2019).

The first CLE peptide was identified for its role in maintaining the stem cell population of the shoot apical meristem; a process during which the peptide AtCLV3 binds to the LRR-RLK AtCLV1 receptor (Clark *et al*., 1995; Fletcher *et al*., 1999). Since then, CLE peptides have been identified for having roles in the root apical meristem (CLE40), vasculature differentiation (TDIF peptides) and inhibition of lateral root development (Hobe *et al*., 2003; Ito *et al*., 2006; Araya *et al*., 2014). A number of CLE peptides are also reported to respond to environmental factors including rhizobia, mycorrhiza, nitrate and phosphate, making them essential for regulating critical plant-microbe interactions as well as macronutrient homeostasis and acquisition (Okamoto *et al*., 2009; Mortier *et al*., 2010; Funayama-Noguchi *et al*., 2011; Reid *et al*., 2011; Araya *et al*., 2014; de Bang *et al*., 2017; Karlo *et al*., 2020).

Nodulation-suppressing CLE peptides are expressed when soil nitrogen availability is high and/or following rhizobia inoculation. They are unique to legumes and have been characterised across different model plants including *Medicago truncatula, L. japonicus*, common bean (*Phaseolus vulgaris*) and soybean (*Glycine max*) (Okamoto *et al*., 2009; Mortier *et al*., 2010; Reid *et al*., 2011; Lim *et al*., 2011; Ferguson *et al*., 2014; Lim *et al*., 2014; Nishida *et al*., 2016). The rhizobia-responsive CLE peptides (M*tCLE12* and *MtCLE13, GmRIC1* and *GmRIC2, PvRIC1* and *PvRIC2*, and *LjCLE-RS1, LjCLE-RS2, LjCLE-RS3*) are induced in the root following inoculation, and their overexpression results in the inhibition of nodule numbers.

To date, nitrate-responsive nodulation-suppressing CLE peptides have not been identified in *M. truncatula* as they have for soybean (GmNIC1a and GmNIC1b), common bean (PvNIC1) and *L. japonicus* (LjCLE-RS2, LjCLE-RS3 and LjCLE40). Bioinformatic analyses by Hastwell *et al*. (2017) identified a total of 52 CLE peptide encoding genes within the *M. truncatula* genome, with the uncharacterised MtCLE34 and MtCLE35 grouping with the previously identified MtCLE12 and MtCLE13 and similar rhizobia- and nitrate-responsive CLE peptides in other model legumes (Hastwell *et al*., 2015b; 2017). Our findings reported here have established new roles for the *M. truncatula* CLE peptides MtCLE34 and MtCLE35 in nitrogen- and rhizobia-signalling during nodulation control.

## Material and Methods

### Plant and bacterial growth conditions

Seeds of wild-type *Medicago truncatula* Jemalong A17 were scarified in 98% sulphuric acid and surface sterilised using 6% (w/v) sodium hypochlorite followed by extensive rinsing in sterile H_2_O. Seedlings were subsequently grown on agar plates containing nitrogen-free Fåhraeus medium (Fåhraeus, 1957) in controlled growth chamber conditions with a photoperiod of 16 h/8 h at 23 °C. Inoculation with *Sinorhizobium meliloti* 1021 (OD_600_ = 0.05) was staggered so that all different time points could be harvested at the same time. Solid Bergersen’s Modified Medium (BMM) was used to culture rhizobia at 28 °C for 2 days and single colonies were grown overnight in liquid BMM before seedling inoculation (Gresshoff *et al*., 1977). The nodulation-susceptible root zone was harvested at 0, 1, 2, 3, 5 and 7 days post inoculation (dpi).

For nitrate treatments, wild-type plants were grown in a sand:perlite (1:2) mixture in controlled growth chamber conditions with a 16 h/8 h photoperiod at 24 °C and 21 °C respectively. Plants were watered as required with nitrogen-limiting Fåhraeus medium (Fåhraeus, 1957) for the first week and supplemented with 0 mM, 2 mM or 10 mM of nitrate (KNO_3_) every second day in the week before sampling the total root.

### Gene expression analysis

Entire roots of 14 day-old nitrate-treated plants or the nodulation-susceptible zone of 11-day old inoculated plants were harvested, snap-frozen and homogenised in liquid nitrogen. The total RNA was extracted using the Maxwell® LEV Simply RNA Tissue kit (Promega) or the RNeasy Plant Mini Kit (Qiagen) according to the manufacturer’s protocol. The quality and quantity of each RNA sample were assessed using the NanoDrop One Spectrophotometer™ (Thermo Fischer Scientific). cDNA was generated using SuperScript® III Reverse Transcriptase (Invitrogen) from 500 ng of DNase-treated RNA. RT-qPCR analysis was conducted using the Roche LightCycler® 96 or the Bio-Rad CFX384 Touch™ with GoTaq SYBR green fluorescence detection (Promega). All reactions were performed in duplicate for at least two biological replicates (n = 11-14) and a target amplicon size of approximately 100 bp. The constitutively expressed Mt40S ribosomal S8 protein was included to normalise expression levels (Mortier *et al*., 2010).

### Bioinformatic analysis

To genetically characterise MtCLE34 and MtCLE35 further, a multiple sequence alignment was performed using Clustal Omega hosted by EMBL-EBI using the prepropeptide amino acid sequences of known nitrate- and rhizobia-responsive CLE peptides in *M. truncatula, L. japonicus*, soybean, common bean, pea and the root-specific CLE peptides of the non-legume *Arabidopsis thaliana* (Goujon *et al*., 2010; Sievers *et al*., 2011; McWilliam *et al*., 2013; Araya *et al*., 2014). In addition, new CLE peptide encoding genes with a potential role in nodulation control were identified using TBLASTN and BLASTN searches against the recently available pea genome (Kreplak *et al*., 2019). A phylogenetic tree was constructed from this alignment using the PhyML plugin in Geneious v.10.0.9, generating a tree based on the maximum likelihood method with 1,000 bootstraps supporting the branches and AtCLV3 as an outgroup (Guindon & Gascuel, 2003). Microsynteny between genomic environments was assessed using Phytozome JBrowse. The orientation, predicted homologues and gene family of five genes directly up- and downstream were evaluated for each gene of interest, consistent with the methods used in Hastwell *et al*. (2015a).

### Sequencing

Sanger sequencing (AGRF) was performed on *M. truncatula* A17 cDNA to confirm premature truncation of the *MtCLE34* transcript. The cDNA was amplified using high-fidelity PrimeSTAR® Max DNA Polymerase (TaKaRa) followed by PCR clean-up. Primers were designed targeting the start codon and a region approximately 100 bp past the predicted ATG-codon to account for signal loss (± 50 bp) during the sequencing process.

### Overexpression and Agrobacterium rhizogenes-mediated transformation

The complete coding sequences of *MtCLE12* (Medtr4g079630), *MtCLE13* (Medtr4g079610) and *MtCLE35* (Medtr2g091125) were cloned into the overexpression vector pCAMBIA1305.1. The PCR products were amplified from genomic DNA using PrimeSTAR® Max DNA Polymerase (TaKaRa). A restriction digest was performed using BglII and BstEII (NEB) followed by ligation using the TaKaRa Mighty Mix ligation kit and heat-shock transformation of competent *Escherichia coli* XL1-Blue cells. Positive clones were confirmed using sequencing. The vectors, including the empty vector, were introduced into the *A. rhizogenes* ARqua1 strain using freeze-thaw transformation (Höfgen & Willmitzer, 1988).

*M. truncatula* seedlings were transformed as described by Boisson-Dernier *et al*. (2001) and hairy roots were selected on modified Fåhraeus medium supplemented with the antibiotic hygromycin (3.5 μg/ml) for two weeks. After that, plantlets were transferred to nitrogen-free Fåhraeus medium and left for 5 days before inoculation with 10 μl *S. meliloti* 1021 (OD = 0.05). Nodules were counted at 10 dpi.

### Statistical analysis

Statistical analyses were performed on all data using the Graphpad Prism v7 software. Following a Shapiro-Wilks normality test, a One-Way ANOVA was used to determine statistically significant differences in relative gene expression of the target genes compared with the respective controls. A Kruskal-Wallis test was used to determine significant differences in nodule number.

## Results

### MtCLE34 and MtCLE35 belong to distinct nodulation-suppressing clades

A phylogenetic tree using the pre-propeptide sequences of the nodulation-suppressing CLE peptides of *M. truncatula, L. japonicus*, soybean, common bean, pea and the non-legume *A. thaliana*, reveals three distinct nodulation-suppressing clades (Figure 1).

**Figure 1.**
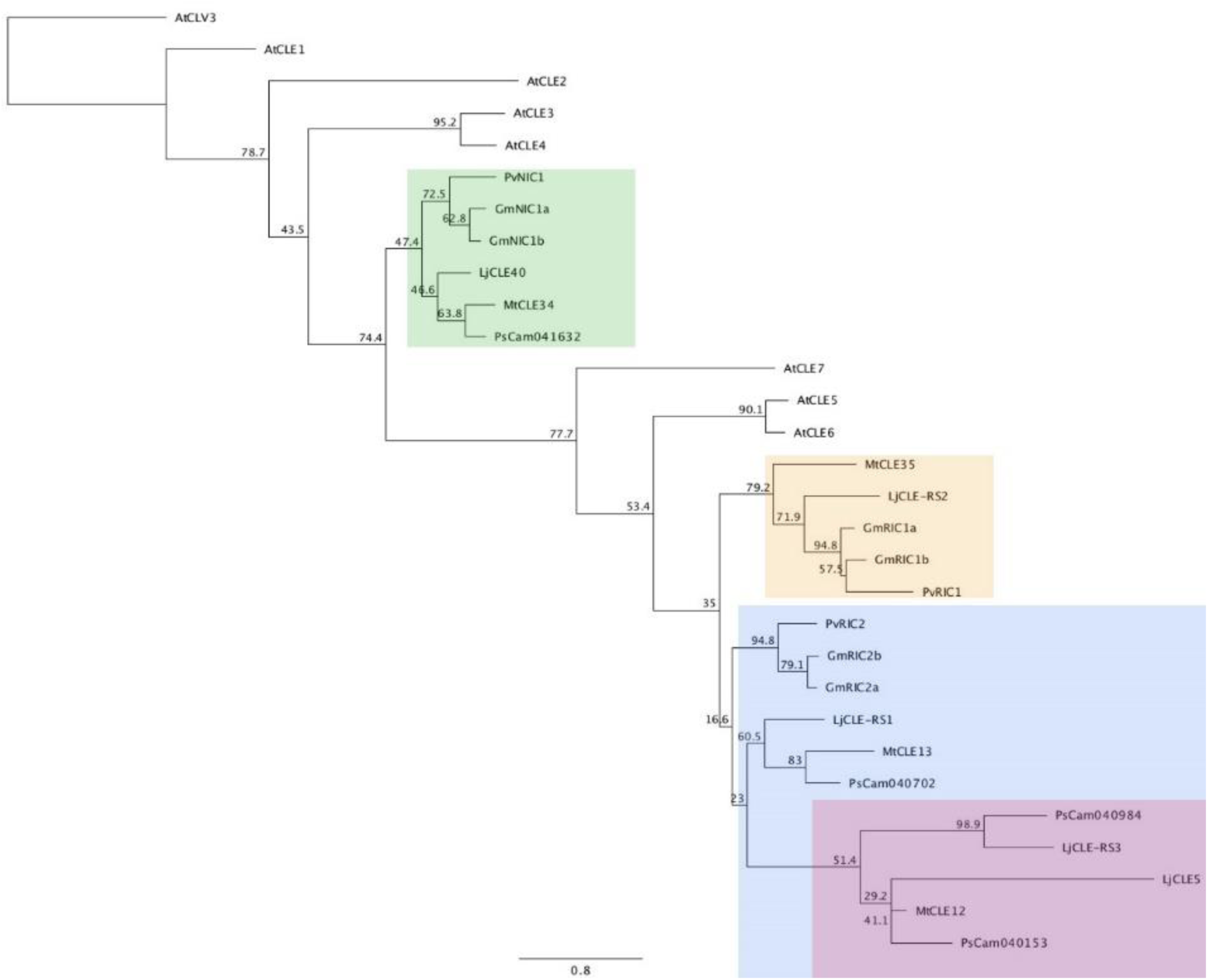
Phylogenetic tree of nitrogen- and/or rhizobia-induced CLE peptides of *Medicago truncatula, Lotus japonicus*, soybean, common bean, pea and *Arabidopsis thaliana*. The full pre-propeptide sequence of each was used. AtCLV3 was added as an outgroup. Three distinct clades (green, orange and blue), with a subgroup (red) were established for the legume sequences. Bootstrap confidence values are shown as percentages from 1,000 bootstrap replications.

MtCLE34 is located in the green group (Figure 1) with the known nitrate-responsive CLE peptides, GmNIC1a/b and LjCLE40. The role of GmNIC1a/b in suppressing nodule organogenesis has been described previously, whereas there has not been conclusive overexpression evidence for the inhibitory role on nodulation of nitrate- and rhizobia-responsive LjCLE40 (Lim *et al*., 2011; Reid *et al*., 2011; Lim *et al*., 2014; Nishida *et al*., 2016). An uncharacterised pea orthologue, PsCam041632, also grouped in this clade indicating a potential role for it in nitrogen-response.

The orange group (Figure 1) contains MtCLE35 as well as rhizobia-responsive GmRIC1a/b, together with LjCLE-RS2 which is induced by both rhizobia and nitrate (Okamoto *et al*., 2009; Lim *et al*., 2011; Reid *et al*., 2011). GmRIC1a/b and LjCLE-RS2 are both widely known suppressors of legume nodulation. Interestingly, a pea orthologue is missing from this group. Since orthologues of genes in the green and orange clades are normally found in tandem pairs, we examined the genomic regions up- and downstream of PsCam041632 from the green group (Figure 2C). Microsynteny of surrounding genes was found to be completely lacking where the pea gene for this group would be expected. Similar investigations using the other nodulation-suppressing pea genes also failed to identify microsynteny. Hence, this region of genes, including the nodulation-suppressing CLE peptide encoding gene, was likely lost through a deletion event.

**Figure 2.**
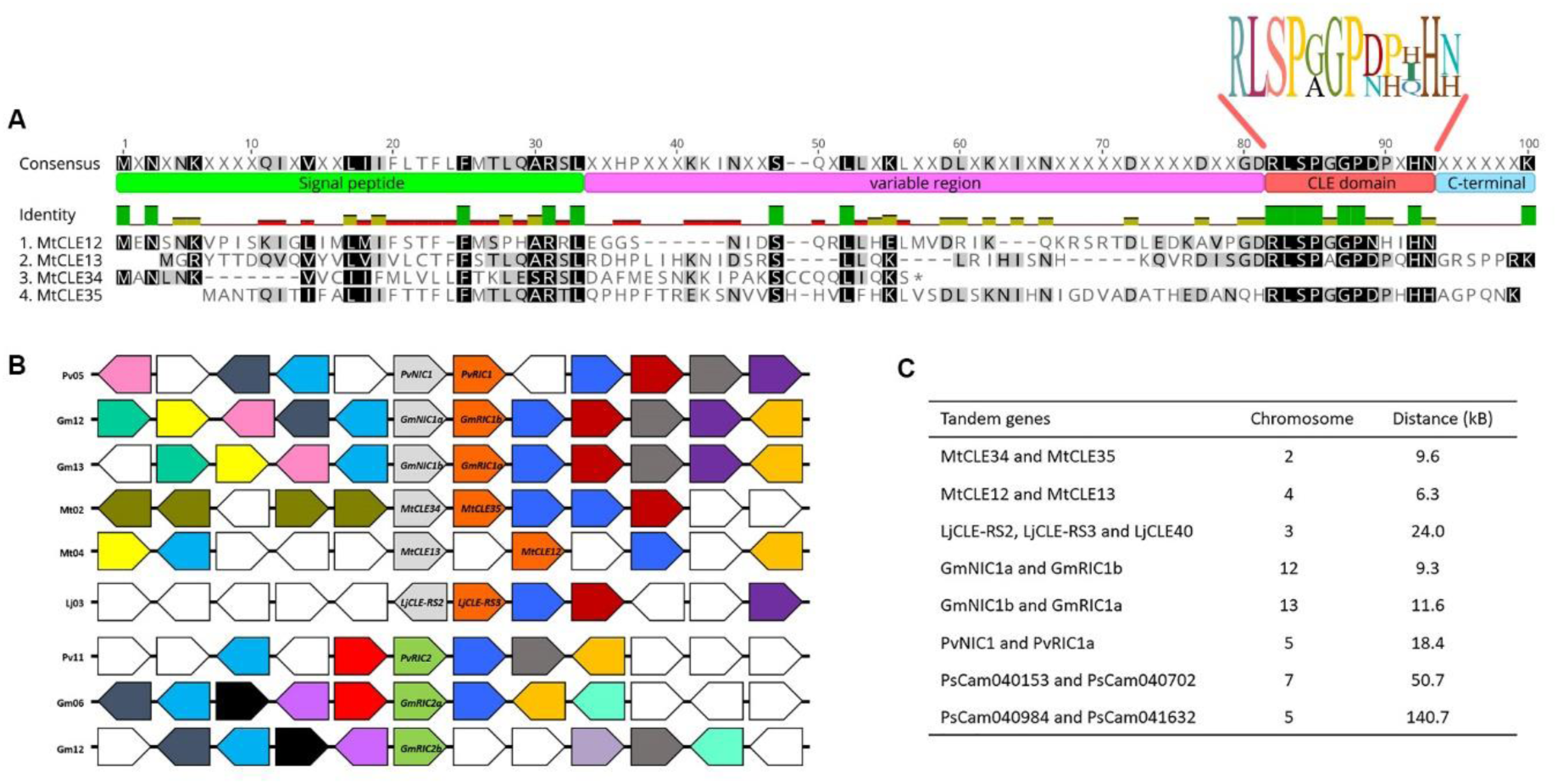
Bioinformatic analysis of nodulation-suppressing CLE orthologues in *Medicago truncatula*, soybean, common bean and *Lotus japonicus*. (A) Shading of amino acids represents conservation among residues with the darker the shading, the higher the level of residue conservation. A sequence logo generated in Geneious v.10.0.9 is shown for the mature CLE peptide. (*) represents a predicted truncation before the mature MtCLE34 peptide. The multiple sequence alignment was generated using Clustal Omega hosted by EMBL-EBI and visualized using Geneious v.10.0.9. (B) Microsynteny of nodulation-suppressive CLE peptides in soybean, common bean, *M. truncatula* and *L. japonicus*. The genes of interest are positioned centrally with arrows indicating the direction of surrounding genes relative to the genes of interest. Genes with a similar putative gene function are shown in the same colours and those with an unrelated or unknown function left uncoloured. (C) Chromosomal location and distance between CLE gene pairs in *M. truncatula, L. japonicus*, soybean, common bean and pea.

MtCLE13 is part of the blue group (Figure 1), which includes a member of every legume species investigated here (GmRIC2a/b, PvRIV2, LjCLE-RS1 and PsCAM040702). Within the blue group is a red subgroup (Figure 1) that includes MtCLE12, LjCLE5, the rhizobia- and nitrate-induced LjCLE-RS3, and two newly identified CLE peptides of pea, PsCam040984 and PsCam040153 (Mortier *et al*., 2010; Nishida *et al*., 2016; Hastwell *et al*., 2017). This subgroup does not contain any orthologues of soybean or common bean.

None of the reported nitrogen-regulated CLE peptides of *A. thaliana* were incorporated into the legume clades established here, which is consistent with previous reports (Figure 1; Hastwell *et al*., 2015a; 2017; 2019). Taken together, this may suggest that the nodulation-suppressing CLE peptides are unique to legumes, and that perhaps the nitrogen-regulated CLE peptides of *A. thaliana* are unique to this species, or to non-legumes in general.

The pre-propeptide sequences of CLE peptides are characterised by a signal peptide domain followed by a variable region and a mature CLE domain at the C-terminal that is highly conserved within gene families and species (Figure 2A; Hastwell *et al*., 2015b). However, MtCLE34 is predicted to be prematurely truncated. Sanger sequencing of *MtCLE34* cDNA confirms the presence of a stop codon (TAG) in the variable region in front of the mature CLE domain. This would prevent the mature CLE peptide ligand from being translated and hence establishes that *MtCLE34* is a pseudogene unable to produce a functional product.

Genomic microsynteny provides further evidence for a possible role of MtCLE34 and MtCLE35 in nodulation control (Figure 2B; 2C). The two genes are located in tandem on chromosome 2 in the *M. truncatula* genome, separated by only 9.6 kb. This suggests a genetic duplication event occurred and that some degree of functional redundancy may exist with MtCLE35 covering for the lack of a functional copy of MtCLE34. This tandem duplication is true for all nodulation-suppressing CLE peptides of the model legumes *M. truncatula, L. japonicus* and soybean suggesting this early duplication occurred before species divergence followed by neofunctionalisation (Figure 2C; Hastwell *et al*., 2015a; 2017). Genomic regions, *i*.*e*., direction and function, are well conserved downstream of the *MtCLE34/MtCLE35* gene pair when compared with *PvNIC1/PvRIC1, GmNIC1a/GmRIC1b, GmNIC1b/GmRIC1a* and *LjCLE-RS2/LjCLE-RS3* (Figure 2B). *MtCLE12* and *MtCLE13* are found on chromosome 4 separated by 6.3 kb with a lower degree of microsynteny compared with other gene pairs.

### MtCLE34 and MtCLE35 are induced by both nitrate and rhizobia

RT-qPCR was used to determine the expression patterns of *MtCLE34* and *MtCLE35* following nitrate treatment or rhizobia inoculation in wild-type roots. Both *MtCLE34* and *MtCLE35* expression were significantly upregulated by the addition of 2 mM or 10 mM KNO_3_ relative to the no-nitrogen control (P ≤ 0.001; Figure 3). Previously, concentrations of 2.5 mM or greater were shown to significantly inhibit nodulation in *M. truncatula*, with the potassium in KNO_3_ having no effect on nodule numbers (van Noorden *et al*., 2016). The transcript abundance of both *MtCLE34* and *MtCLE35* gradually increased as the nitrate concentration increased (Figure 3). This is consistent with previous reports demonstrating a reverse correlation between increasing concentrations of available nitrate and reduced nodule numbers (Carroll *et al*., 1985; Barbulova *et al*., 2007; van Noorden *et al*., 2016). In contrast, expression of the known nodule-suppressing CLE peptide encoding genes, *MtCLE12* and *MtCLE13*, was unaffected by nitrate treatment (*P* ≥ 0.1; Figure 3).

**Figure 3.**
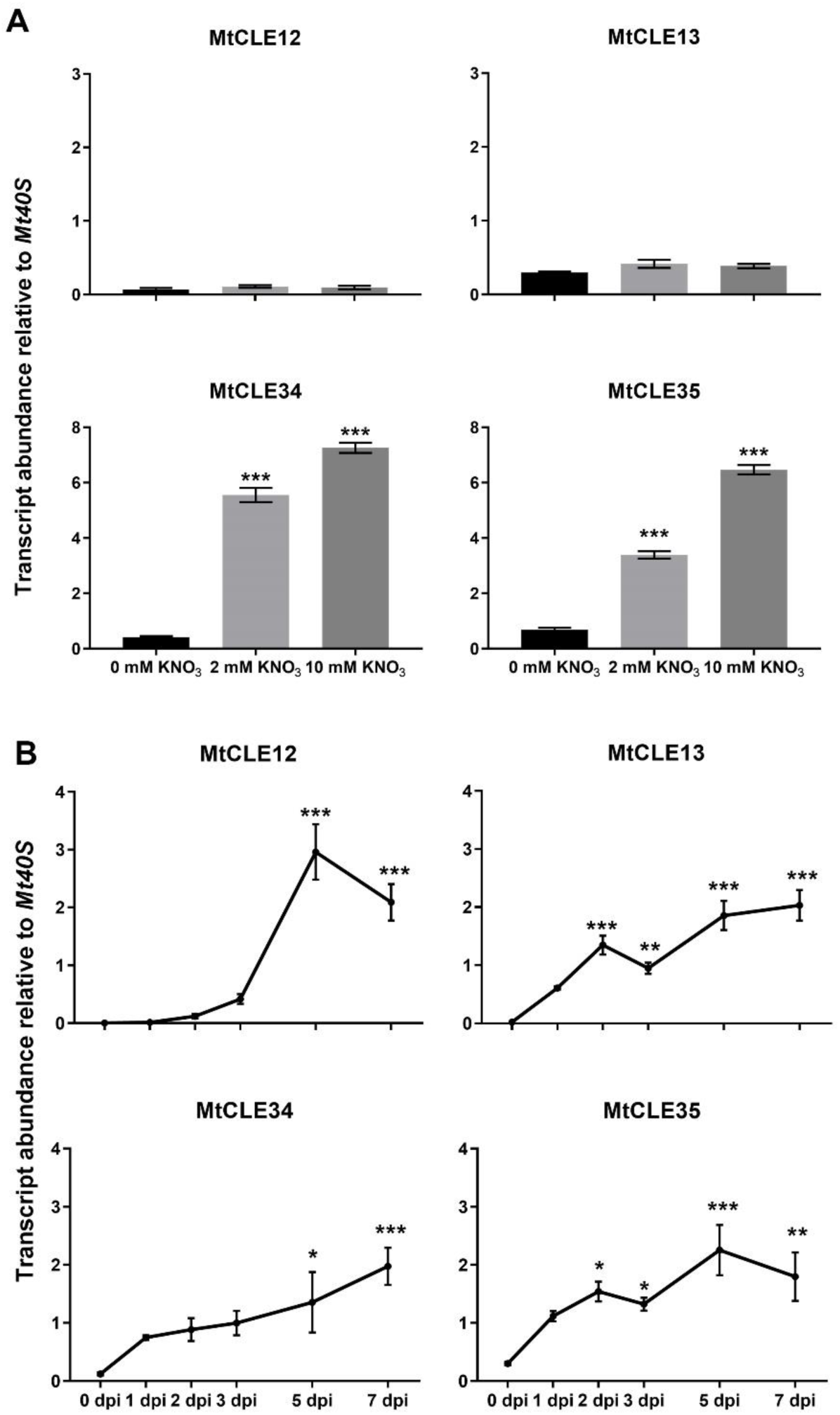
Expression of *MtCLE12, MtCLE13, MtCLE34* and *MtCLE35* in response to (A) nitrate treatment and (B) rhizobia inoculation. (A) Wild-type *M. truncatula* plants (cv. Jemalong A17) were treated with 0 mM, 2 mM or 10 mM of nitrate (KNO_3_), after which entire roots of 14-day old plants were harvested. (B) Following inoculation with *Sinorhizobium meliloti*, the zone of nodulation above the root tip was harvested at 0, 1, 2, 3, 5 and 7 days post inoculation (dpi). Transcript abundance is relative to the housekeeping gene *Mt40s ribosomal protein S8*. Bars represent means ± SEM of at least two biological replicates (n = 11-14 per replicate). Asterisks indicate significant differences in transcript abundance compared to the control (One-Way ANOVA, \*\*\**P* ≤ 0.001, ** *P* ≤ 0.01, \**P* ≤ 0.05).

In addition to nitrogen induction, the rhizobia-responsiveness of *MtCLE12, MtCLE13, MtCLE34* and *MtCLE35* was investigated by inoculating wild-type plants with *S. meliloti*, after which the zone of nodulation was harvested at 0, 1, 2, 3, 5 and 7 dpi. As expected, *MtCLE12* and *MtCLE13* were induced by the bacteria, with a subtle difference in the timing of their onset (Figure 3; Mortier *et al*., 2010). *MtCLE12* was induced at 5 dpi, reaching a peak in transcript abundance after which levels dropped off slightly (*P* ≤ 0.001), while *MtCLE13* was significantly upregulated as early as 2 dpi with their transcript levels increasing up to 7 dpi (*P* ≤ 0.005; Figure 3). The newly identified *MtCLE35* was induced by rhizobia at 2 dpi (*P* ≤ 0.05), in a pattern similar to that of *MtCLE13*. Interestingly, despite *MtCLE34* clustering with the CLE peptides acting in nitrate-dependent regulation of nodulation, it was also induced by rhizobia at 5 and 7 dpi (*P* = 0.02 and *P* ≤ 0.001 respectively; Figure 3).

### MtCLE35 inhibits nodulation

To determine whether MtCLE35 inhibits nodulation, overexpression of *MtCLE35* using *A. rhizogenes*-mediated hairy root transformation in a wild-type (A17) background was performed. *MtCLE12* and *MtCLE13* were also overexpressed. *MtCLE34* overexpression was not assessed because it was confirmed here to be a pseudogene based on sequencing. Overexpression of all three CLE peptides reduced nodule numbers to a similar extent compared with the empty vector control (*P* < 0.001; Figure 4) confirming a role for MtCLE35 in nodulation control.

**Figure 4.**
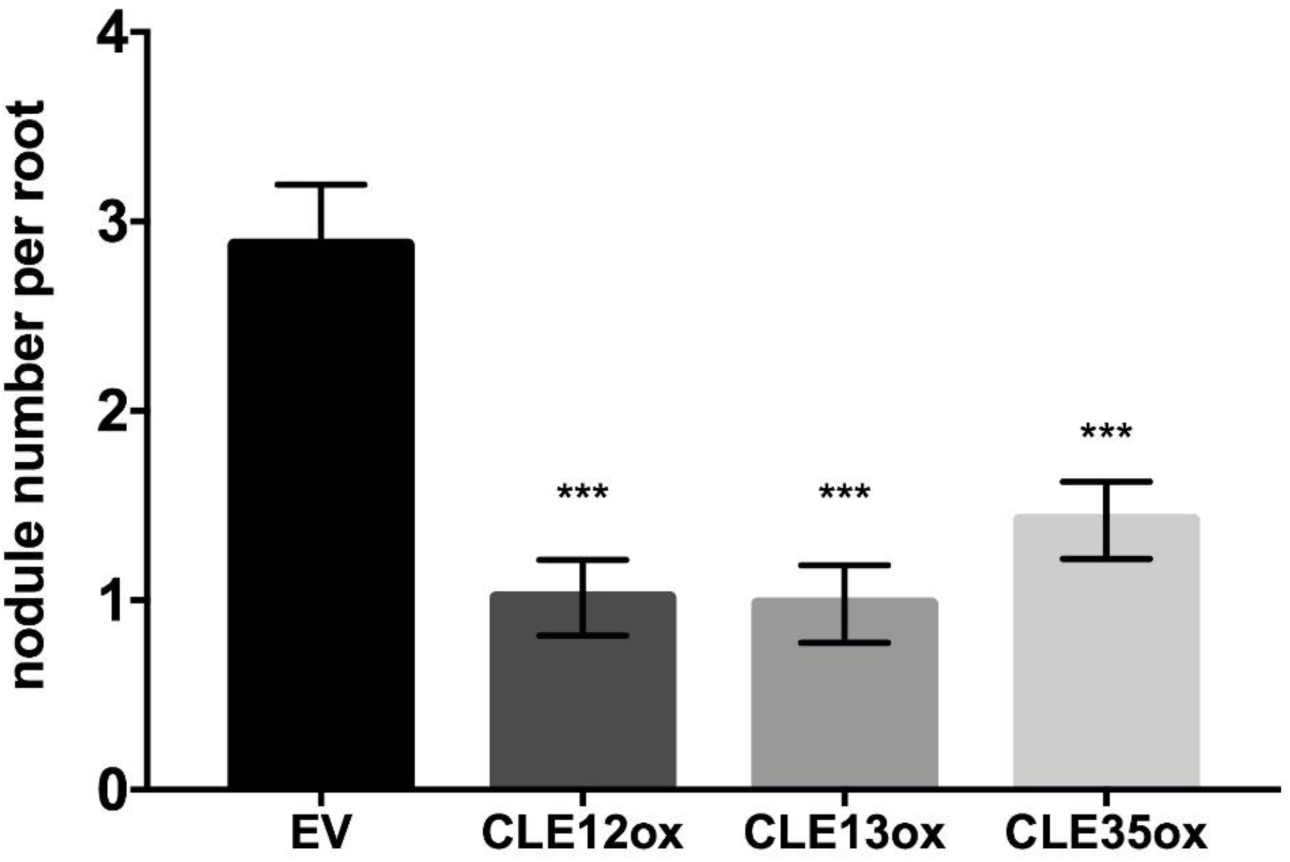
Nodule numbers of wild-type A17 plants overexpressing *MtCLE12, MtCLE13* or *MtCLE35*. Bars represent means ± SEM. Asterisks indicate significant differences in nodule number compared to the empty vector control (n = 48-78; Kruskal-Wallis, *** *P* ≤ 0.001).

## Discussion

Nitrogen homeostasis is crucial for plant development and requires the integration of internal and external signals (*e*.*g*., nutrient status). Legumes tightly regulate their nodule number, size and nitrogen fixation rates in the presence of nitrogen (Streeter, 1988; Ferguson *et al*., 2019). While nitrogen-responsive CLE peptides have been previously identified in soybean, common bean and *L. japonicus*, they had not yet been characterised in *M. truncatula*, with only rhizobia-responsive CLE peptides reported (MtCLE12 and MtCLE13; Mortier *et al*., 2010). Moreover, pea orthologues of the nodulation-suppressing CLE peptides had not yet been thoroughly identified (Hastwell *et al*., 2019).

Our findings indicate that MtCLE12 and MtCLE13 have a distinct role in rhizobia-induced AON, whereas MtCLE35 can regulate nodule numbers in both a rhizobia- and nitrate-induced manner. Previous studies have shown that triarabinosylation of the peptides is required for SUNN-dependent nodule control (Schnabel *et al*., 2011; Kassaw *et al*., 2017; Imin *et al*., 2018). MtCLE34 likely had a similar role to MtCLE35, but the gene encoding it has been lost through a naturally-occurring mutation and a lack of evolutionary pressure that resulted in pseudogenisation and loss of the peptide ligand. Interestingly, *L. japonicus* also contains a mis-annotated pseudogene (*LjCLE5*) that is orthologous to *MtCLE12* (Hastwell *et al*., 2017), although it is reported to not be expressed upon either rhizobia inoculation or nitrate treatment (Okamoto *et al*., 2009).

The nodulation-suppressing CLE peptides of legumes can be categorised into three distinct clades. The green clade contains MtCLE34, GmNIC1a/b, PvNIC1, PsCam041632 and LjCLE40. Indeed, all species explored here have a peptide in this group. They are induced by nitrate, with *MtCLE34* and *LjCLE40* also induced by rhizobia (Nishida *et al*., 2016).

The orange group contains MtCLE35, GmRIC1a/b, PvRIC1 and LjCLE-RS2. All of the genes encoding these peptides are induced by rhizobia, with *MtCLE35* and *LjCLE-RS2* also induced by nitrate (Okamoto *et al*., 2009; Reid *et al*., 2011; Nishida *et al*., 2016). Interestingly, pea seems to have lost its copy based on our bioinformatic analyses and genetic synteny investigations in the regions surrounding the other four CLE peptide genes of pea reported here.

The blue clade of nodulation-suppressing CLE peptides contains members that are only induced by rhizobia (*i*.*e*., MtCLE13, GmRIC2a/b, PvRIC2, and LjCLE-RS1, as well as PsCam040702). Through duplication events, this group led to the red subgroup that lack soybean and common bean orthologues (Okamoto *et al*., 2009; Mortier *et al*., 2010; Reid *et al*., 2011). The lack of these orthologues suggests that the red subgroup was either lost in soybean and bean, or originated in pea, *M. truncatula* and *L. japonicus* after the Phaseoloid clade diverged from the Hologaligena clade >54 million years ago (Lavin *et al*., 2005; Egan & Doyle, 2010; Vanneste *et al*., 2014).

Of the red subgroup, one branch contains MtCLE12, LjCLE5, and PsCam040153. The second contains LjCLE-RS3 and PsCam040984. Unlike the other members investigated to date, *LjCLE-RS3* is the only one to also be induced by nitrate, but its induction is low compared to *LjCLE-RS2* (Nishida *et al*., 2016). The missing *M. truncatula* CLE peptide in the second branch may have been lost, which is common for duplicated genes over time (Lynch & Conery, 2000). Alternatively, it could be located in a part of the genome that has not been fully sequenced, but this seems unlikely as the *M. truncatula* genome has been thoroughly re-sequenced multiple times (Mt4.0v1; Tang *et al*., 2014).

Each of the legume species investigated here has CLE peptides in multiple clades. Their genes are often found directly adjacent on chromosomes, indicating they arose from tandem duplication(s) before speciation, after which neofunctionalisation occurred. These legume-specific peptides most likely evolved from existing CLE peptides acting in other developmental processes.

A role for small secreted peptides in macronutrient response was previously investigated by de Bang *et al*. (2017). In this study, *MtCLE34* was downregulated in response to deficiencies in nitrogen, as well as phosphorus and sulphur, then was strongly upregulated following the re-supplementation of these macronutrients (de Bang *et al*., 2017). This indicates a more general role for MtCLE34 might have existed prior to it accumulating a null mutation. It is possible that MtCLE35 also responds to a number of macronutrients to regulate general root plasticity and nutrient homeostasis.

In *A. thaliana*, a number of CLE peptides have been identified that respond to nitrate. However, none of these CLE peptides group with the legume CLE peptides (Hastwell *et al*., 2017) and they conversely are upregulated following nitrogen deficiency (*AtCLE1/3/4/7*; Araya *et al*., 2014). Overexpression of these four peptides results in an inhibition of lateral root development (*i*.*e*., lateral root emergence), while the *Atclv1* receptor mutant is characterised by increased lateral root length and density, indicating these peptides function in nitrogen foraging (Araya *et al*., 2014). Similarly, AON LRR-RLK mutants often have aberrant root phenotypes that include shorter primary roots and higher lateral root density (*e*.*g*., Wopereis *et al*., 2000; Schnabel *et al*., 2005). In addition, some root architecture responses to nitrate are differentially affected in the *M. truncatula* AON mutants (Goh *et al*., 2018).

Unlike *MtCLE34* and *MtCLE35*, which respond to both rhizobia and nitrate, *MtCLE12* and *MtCLE13* are upregulated by rhizobia but not by nitrate treatment. MtCLE13 groups closely with the rhizobia-responsive CLE peptides GmRIC2a/b and LjCLE-RS1. MtCLE12 forms a clade with LjCLE-RS3, which seems to act differently than the other nodulation-suppressing CLE peptides. Indeed, both *MtCLE12* and *LjCLE-RS3* are upregulated at a later stage of nodulation with the latter also being induced in the presence of nitrogen and rhizobia. *LjCLE-RS3* overexpression also results in a weaker inhibition of nodulation compared with *LjCLE-RS1* and *LjCLE-RS2* (Nishida *et al*., 2016). While MtCLE13 is nodule-specific, MtCLE12 expression also occurs in the root tip, cotyledons and first leaves (Mortier *et al*., 2010). In addition, it is not induced by exogenous application of cytokinin, unlike *MtCLE13* (Mortier *et al*., 2010). Both cytokinin and the consequent upregulation of NODULE INCEPTION (NIN), are crucial for nodule primordia establishment and infection (Murray *et al*., 2007; Tirichine *et al*., 2007; Plet *et al*., 2011). NIN has also been shown to bind to the promoter region of these CLE peptides linking it to AON and nitrate-dependent control of nodulation (Soyano *et al*., 2014; 2015; Lin *et al*., 2018; Nishida *et al*., 2018).

Maintaining multiple CLE peptides able to regulate nodule numbers must be important for effective control of nodulation and hence why legume species have retained so many throughout evolution. Subtle temporal differences in the expression of the nodulation-suppressing CLE peptides indicates that timing of expression is also crucial. More work is now needed to understand the subtle and sophisticated roles each of these peptide hormones play in nodulation control, and possibly also in other aspects of plant development and nutrient response.

## Author contributions

BF devised the project and BF and PM supervised. CM and AH performed bioinformatic analyses. HS assisted with cloning and UM conducted the overexpression experiment. CM performed gene expression analysis, analysed the data, prepared figures and wrote the manuscript with BF and additional input from all authors. All authors read and approved the manuscript.

## Acknowledgements

This work was funded by the Australian Research Council (ARC) Grants DP130102266, DP130103084 and DP190102996; and the Hermon Slade Foundation. CM is supported by an RTP Scholarship. AHH is funded by an ARC Discovery Early Career Research Award (DE200100800) and the Molly-Budtz Olsen Fellowship from The Fellowship Fund Inc. We would like to acknowledge Bowen Harding and Jason Ng for their technical assistance.

